# Prefrontal allopregnanolone mediates the adverse effects of acute stress in a mouse model of tic pathophysiology

**DOI:** 10.1101/2022.10.29.514372

**Authors:** Roberto Cadeddu, Meghan Van Zandt, Karen Odeh, Collin J Anderson, Deirdre Flanagan, Peter Nordkild, Christopher J Pittenger, Marco Bortolato

## Abstract

Ample evidence suggests that acute stress can worsen symptom severity in Tourette syndrome (TS); however, the neurobiological underpinnings of this phenomenon remain poorly understood. We previously showed that acute stress exacerbates tic-like and other TS-associated responses via the neurosteroid allopregnanolone (AP) in an animal model of repetitive behavioral pathology. To verify the relevance of this mechanism to tic pathophysiology, here we tested the effects of AP in a mouse model recapitulating the partial depletion of cholinergic interneurons (CINs) in the striatum seen in *postmortem* studies of TS. Mice underwent targeted depletion of striatal CINs during adolescence and were tested in young adulthood. Compared with controls, CIN-depleted male mice exhibited several TS-relevant abnormalities, including deficient prepulse inhibition (PPI) and increased grooming stereotypies after a 30-min session of spatial confinement, a mild acute stressor that increases AP synthesis in the prefrontal cortex (PFC). These effects were not seen in females. Systemic and intra-PFC AP administration dose-dependently worsened grooming stereotypies and PPI deficits in CIN-depleted males. Conversely, both AP synthesis inhibition and pharmacological antagonism reduced the effects of stress. These results further suggest that AP in the PFC mediates the adverse effects of stress on the severity of tics and other TS manifestations. Future studies will be necessary to confirm these mechanisms in patients and define the circuitry responsible for the effects of AP on tics.

## INTRODUCTION

Tics are sudden, non-rhythmic, stereotypical movements or utterances characterized by variable intensity and complexity. The most disabling tic disorder, Tourette syndrome (TS), is a childhood-onset condition characterized by multiple motor tics and at least one phonic tic for over one year^1^. TS is complicated by the high prevalence of comorbid psychiatric disorders, such as obsessive-compulsive disorder (OCD), attention-deficit hyperactivity disorder (ADHD), anxiety, and depression^2,3^. Current pharmacotherapies for TS remain unsatisfactory. The main pharmacological treatments for TS, dopaminergic antagonists and α_2_ adrenergic agonists^4^, are associated with inconsistent efficacy and significant adverse effects^5–7^.

Tics fluctuate in number, frequency, intensity, and complexity^8^. While the biological causes of these fluctuations remain elusive, several lines of evidence point to environmental stress as a crucial influence on tic severity. For example, tic severity is associated with stressful life events^9,10^. This relationship has been confirmed by longitudinal analyses, which have documented that cumulative psychosocial stress predicts future tic severity^11^. Furthermore, tic severity is correlated with self-report ratings of daily stress^12^ and recent adverse events^13^. Tics can be exacerbated by specific stressors, such as hypostimulation and fatigue^14^; the contribution of acute psychosocial stress may be more complex^15^.

Tic disorders are associated with anatomical and functional alterations within the cortico-striatal-thalamo-cortical circuitry. The causes of these alterations remain poorly understood. Several studies have documented a selective loss in cholinergic and parvalbumin-positive GABAergic interneurons in the dorsal striatum of individuals with severe TS^16–18^. It is possible that a local reduction in striatal interneurons (perhaps due to genetic and early-life inflammatory factors) may lead to the formation of “focal disinhibition areas” in the striatum^19^. Abnormal activation of these foci may lead to the involuntary execution of tics.

Building on these observations, Xu *et al*.^20^ modeled the loss of striatal cholinergic interneurons (CIN) through a targeted ablation in the dorsal striatum of adult mice. This manipulation produced no overt baseline behavioral alterations, but it predisposed mice to exhibit grooming stereotypies in response to acute stress^20^. Subsequent work revealed a sexual dimorphism to this effect, with stereotypies emerging only in males, not in identically manipulated females^21^. The lack of baseline behavioral deficits in these mice may partially reflect the fact that CINs were depleted in adulthood following normal development; this may have missed critical neurodevelopmental components. On the other hand, the stress sensitivity seen in these animals is strikingly reminiscent of the worsening of tics and other symptoms after stress in TS patients. The mechanisms underlying these relationships remain unclear.

Growing evidence suggests that the pathophysiology of TS may be modulated by neurosteroids^22^. These mediators play a crucial role in modulating the stress response^23–25^. Using a different mouse model of TS with high face and predictive validity, the D1CT-7 mouse, we have shown that stress exacerbates tic-like behaviors as well as other TS-relevant effects^26^ through the action of the neurosteroid allopregnanolone (AP; 3α,5α-tetrahydroprogesterone)^27^. This metabolite of progesterone is increased in response to acute stress and activates GABA_A_ receptors^28,29^. AP elicited tic-like movements, increased stereotyped behavior, and induced PPI deficits in D1CT-7 mice^27^. Furthermore, isoallopregnanolone (isoAP), the 3β-epimer of AP that acts as an antagonist to the AP-binding site within GABA_A_ receptors, countered the adverse effects of stress on tic-like behaviors and PPI deficits in this model^30^.

The present study was designed to model the impact of early-life CIN depletion in mice and assess its implications for sensorimotor gating and stress susceptibility. To test the effects of AP in this model, we tested whether this neurosteroid can aggravate tic-like stereotypies and whether antagonizing its actions can block the effects of acute stressors. This extends our previous work in the D1CT-7 mouse, which has limited construct validity, to a model of TS pathophysiology that recapitulates a neuropathological abnormality associated with refractory tics^17^.

## MATERIALS AND METHODS

### Animals

Homozygous choline acetyltransferase (ChAT)-*cre* female mice (B6; 129S6-ChAT^tm2(cre)Lowl/J^ obtained from The Jackson Laboratory, 006410) were crossed with C57BL/6J male mice (The Jackson Laboratory, 000664; Bar Harbor, ME). Male and female hemizygous ChAT-*cre* transgenic mice were produced in our animal facilities by crossing hemizygous male ChAT^tm2(cre)Lowl/J^ mice with female wildtype C57BL/6J mice or hemizygous female ChAT^tm2(cre)Lowl/J^ mice with male wildtype C57BL/6J mice. Genotypes were confirmed by PCR. Experiments were performed using adult male and female mice (3-4-month-old). All animals were allowed to acclimate to facilities for 7-10 days before behavioral testing. Housing facilities were maintained at 22°C with a reversed 12-hour light/dark cycle (lights off at 6:00 AM, lights on at 6:00 PM). Experimental manipulations were carried out during the dark cycle (between 9:00 AM and 5:00 PM). Every effort was made to minimize the number and suffering of animals. Thus, animal numbers for each experiment were determined through power analyses conducted on preliminary results. All experimental procedures were approved by the local Institutional Animal Care and Use Committees.

### CIN depletion procedure

CIN depletion was induced in ChAT-cre hemizygous mice using a modified version of the protocol described by Xu *et al.^20^*, specifically designed to enable CIN depletion in early developmental stages. Male and female pups at postnatal day (PND) 4 underwent bilateral stereotaxic infusion of either a viral construct harboring the simian diphtheria toxin receptor (sDTR; A06) or its control (C06)^20^. Briefly, after anesthesia was induced in a pup by hypothermia, its head was gently placed between the ear bars of a neonatal stereotaxic frame (Stoelting; Wood Dale, IL). The following coordinates were used to target the striatum: AP - 0.5 mm; ML ± 1.6 mm; DV - 2.8 mm. A 5 μl Hamilton Neuros syringe (Reno, NV) was used to inject 0.25 μl of viral construct or control vector bilaterally. The syringe was left in place for at least 2 minutes after completion of the infusion to eliminate backflow. The pup was then placed on a warming pad until its body temperature and skin color returned to normal, and afterward, it was returned to its mother. Following surgery, mice were left undisturbed in their home cage until PND 18, when they received a single intraperitoneal (IP) injection of DT (Sigma, St. Louis, MO) at a dose of 1 μg/kg. Mice were weaned at PND 21 and group-housed in same-sex sibling pairs until experimental procedures.

### Drugs

AP (Tocris Bio-Techne, Minneapolis, MN) and finasteride (AstaTech, Bristol, PA) were dissolved in 2.5% DMSO, 2.5% Tween 80, and 0.9% NaCl. The AP 3β-epimer isoAP (3β,5α, tetrahydroprogesterone) was donated by Asarina Pharma AB (Solna, Sweden). IsoAP was suspended in 3% hydroxypropyl β-cyclodextrin and administered IP. For regional intracerebral infusions, AP was dissolved in a Tween 80/Ringer solution (final concentration, 1:1, v:v); AP was then infused in a cyclodextrin/Ringer solution (final concentration 1:5, v:v). For regional intracerebral studies, isoAP was infused in Ringer solution (final concentration 1:1, v:v). The D_1_ receptor (D_1_R) antagonist SCH23390, the D_2_ receptor (D_2_R) antagonist haloperidol, and the α_2_-adrenergic agonist clonidine (Sigma, St. Louis, MO) were dissolved in 0.9% NaCl and administered IP. The injection volume for all systemic administrations was 10 ml/kg.

### Stereotaxic surgery for regional intracerebral studies

CIN-depleted and control mice were deeply anesthetized with xylazine/ketamine (20‧80 mg‧kg^-1^, IP) and then placed into a stereotaxic frame (David Kopf Instruments, Tujunga, CA). The target locations for cannulation were, from bregma: i) medial prefrontal cortex (mPFC): AP + 1.8 mm, ML ± 0.3 mm, DV – 2.5 mm from the skull surface; ii) dorsolateral striatum (DLS): AP + 0.5 mm, ML ± 2.2 mm, DV – 2.2 mm, according to the coordinates of Paxinos and Watson^35^. Mice were allowed to recover in their home cages (single-housed), and after one week, they received bilateral microinjections in the targeted area (according to the specific experimental design). Microinjections were performed by gently restraining the mouse, removing the stylet, and replacing it with the injector (P1 Technologies Inc.) connected by a 250 μl Hamilton syringe via PE tethers. A microinfusion pump (KD Scientific, Holliston, MA) delivered 0.5 μl/side of drug solution (or its vehicle solution) at a constant flow/rate of 0.25 μl/min. After infusion, the injector was left in place for 2 minutes to allow fluid diffusion. After behavioral tests, animals were sacrificed, and the histological verification of the cannula location was confirmed. Animals with errant locations of either device or damage to the targeted area(s) were excluded from the analysis.

### Spontaneous and stress-induced grooming behavior

Self-grooming behavior of freely moving mice was assessed in a regular mouse cage (29 × 17 × 13 cm) with bedding material. After a 10-min familiarization period, the behavior was video-recorded for 30 min. To test whether grooming behavior may be affected by a mild acute stressor, we used spatial confinement^26^. Briefly, animals were confined within a clear, bottomless Plexiglas cylinder (10□cm in diameter × 30□cm in height), which was placed in a familiar cage, and deeply sunk into the bedding to ensure stability. The procedure lasted 40 minutes, and behaviors were video-recorded for the last 30 min to avoid potential behavioral alterations caused by neophobia induced by the exposure to the unfamiliar enclosure. Grooming behavior was scored by trained observers blinded to genotype and treatment and included complete and incomplete sequences of licking, scratching, and washing the paws, head, body, and tail. To avoid potential carry-over effects of stress, each animal was used only once in this procedure.

### Prepulse inhibition of startle reflex

Startle testing was conducted in sound-attenuating ventilated startle chambers (SR-LAB, San Diego Instruments, San Diego, CA) as previously detailed^31^. Two different protocols were used: 1) acoustic PPI and 2) tactile-acoustic PPI (with tactile pulses and acoustic prepulses). The acoustic PPI protocol featured a 70 dB background white noise (5 minutes acclimation period), followed by three consecutive blocks of “pulse”, “prepulse + pulse”, and “no stimulus” trials. During the first and third blocks, mice received five “pulse alone” trials of 115 dB. During the second block, animals underwent a pseudo-random sequence of fifty trials, consisting of “pulse-alone” trials (twelve events), pulses preceded by 73, 76, or 82 dB “prepulses” (ten events for each level of prepulse), and “no stimulus” trials (eight events, consisting of background white noise delivery). The mixed PPI protocol followed the same sequence, but pulse-alone stimuli were delivered via a 2-mm plastic tube used to deliver air-puff stimuli, was projected directly downward towards the animal’s back. The source of air-puff stimulation was compressed air delivered at a constant pressure of 15 psi. The resulting “air-puff” then stimulated the dorsal torso of the animals through a hole in the top of the enclosure. The animal’s average startle amplitude in response to bursts of air-puff stimulation was recorded and analyzed. Startle-Only trials consisted of a burst of air-puff stimulation lasting 40 ms in duration. Prepulses in the tactile-acoustic PPI protocol consisted of auditory stimuli identical to those used in the auditory PPI protocol. Intertrial intervals (ITI) in both protocols were selected randomly between 10 and 15 seconds. A dynamic calibration system was used to ensure comparable sensitivities across chambers. Percent PPI (%PPI) was calculated using the formula (Mean Startle Amplitude, MSA):

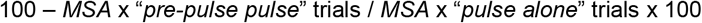

### Assessment of eyeblinks and fine head movements

Eye blinking and head movements were monitored during spatial confinement (as described above). The cylinder was mounted on a square platform adjacent to four video cameras placed on each side^32^. This configuration allows experimenters to monitor eyeblinks remotely and continuously, in a non-invasive fashion, and without the employment of head restraint bars. Animals were exposed to the apparatus for 2 days before testing. Eyeblinks and axial head jerks were scored by blinded observers.

### Locomotor activity in the open field

Horizontal locomotor activity was tested in a black Plexiglass open field arena (40 × 40 × 40 cm), as previously described^33^. Briefly, mice were placed in the center of the arena and allowed to explore freely for 5 minutes. Locomotor activity and time in the center (defined as a central 28.3 × 28.3 cm square) were scored and analyzed using behavioral tracking software (EthoVision XT, Noldus, Wageningen, The Netherlands).

### Marble-burying test

Each mouse was transferred individually into a clean home cage (29 × 17 × 13 cm) containing 16 colored marbles homogenously distributed over 5 cm bedding for a 15-minute trial. Testing was performed with normal (100 lux) and high illumination (1000 lux) conditions. Buried marbles were counted blind to condition and treatment before starting the procedure and at the end of testing.

### Elevated plus maze

Anxiety-like behavior was assessed using an elevated plus-maze as described previously^34^. The maze consisted of two open arms (35 × 6 cm) and two closed arms (35 × 6 × 21 cm) extending from a central platform (6 × 6 cm) elevated at 74 cm from the ground. Mice were placed in the center area of the maze with their head directed toward a closed arm. The time spent in the open arms (with all four paws in the arm) was measured and expressed as a percentage of the total time in the maze. The number of entries and head dips were also monitored. The overall duration of the test was 5 minutes.

### Immunohistochemistry and Quantification

Mice were sacrificed and perfused with 4% paraformaldehyde, then the brains were removed and fixed overnight, followed by cryoprotection in 30% sucrose. The brains were then frozen in NEG-50 (Richard-Allan Scientific) tissue freezing medium and sectioned on a Leica cryostat at 30 μm thick and mounted on glass slides. Slides with brain sections containing the dorsolateral striatum were incubated in primary antibodies overnight (22-26 hours) – mouse anti-ChAT (Thermo Fischer, MA5-31383) and chicken anti-GFP (Aves, GFP-1020). Sections were then incubated for two hours in secondary antibodies – goat anti-mouse 568 (Thermo Fischer, A-11031) and goat anti-chicken 488 (Thermo Fisher, A-11039), followed by coverslipping with ProLong Gold with DAPI for nuclear labeling (Thermo Fischer, 36931). Sequential images of the dorsal striatum were taken on a Zeiss Scope.A1 using a Plan-APOCHROMAT 10x objective and an Axiocam 503 mono. These images were then stitched together using the Stitching Plugin in FIJI (FIJI is Just ImageJ, Schindelin *et al.^36^)* using the default values on the Sequential Images setting. Cell counting was then performed with counters blind to sex and treatment of animal in these reconstructed sections.

### Statistical analyses

Normality and homoscedasticity of data distribution were verified using Kolmogorov-Smirnov and Bartlett’s tests. Statistical analyses of parametric data were performed using t-test, one-way ANOVA or two-way ANOVA, as appropriate. *Post-hoc* comparisons were performed by Tukey’s test. Statistical analyses of nonparametric data for two-group comparisons were performed by Mann-Whitney test. Significance thresholds were set at 0.05.

## RESULTS

### Early-life CIN depletion leads to TS-related behavioral alterations in male mice

To evaluate the efficacy of CIN depletion, we counted ChAT-positive neurons in the striatum of mice treated with an active virus (A06) compared to counterparts treated with the control virus (C06). Because our past work showed sexual dimorphism in the effects of interneuron depletion, we examined male and female mice separately. In males, DT administration successfully reduced the number of inhibitory interneurons in the striatum in A06-treated mice (Fig. 1A-B).

**Fig. 1.**
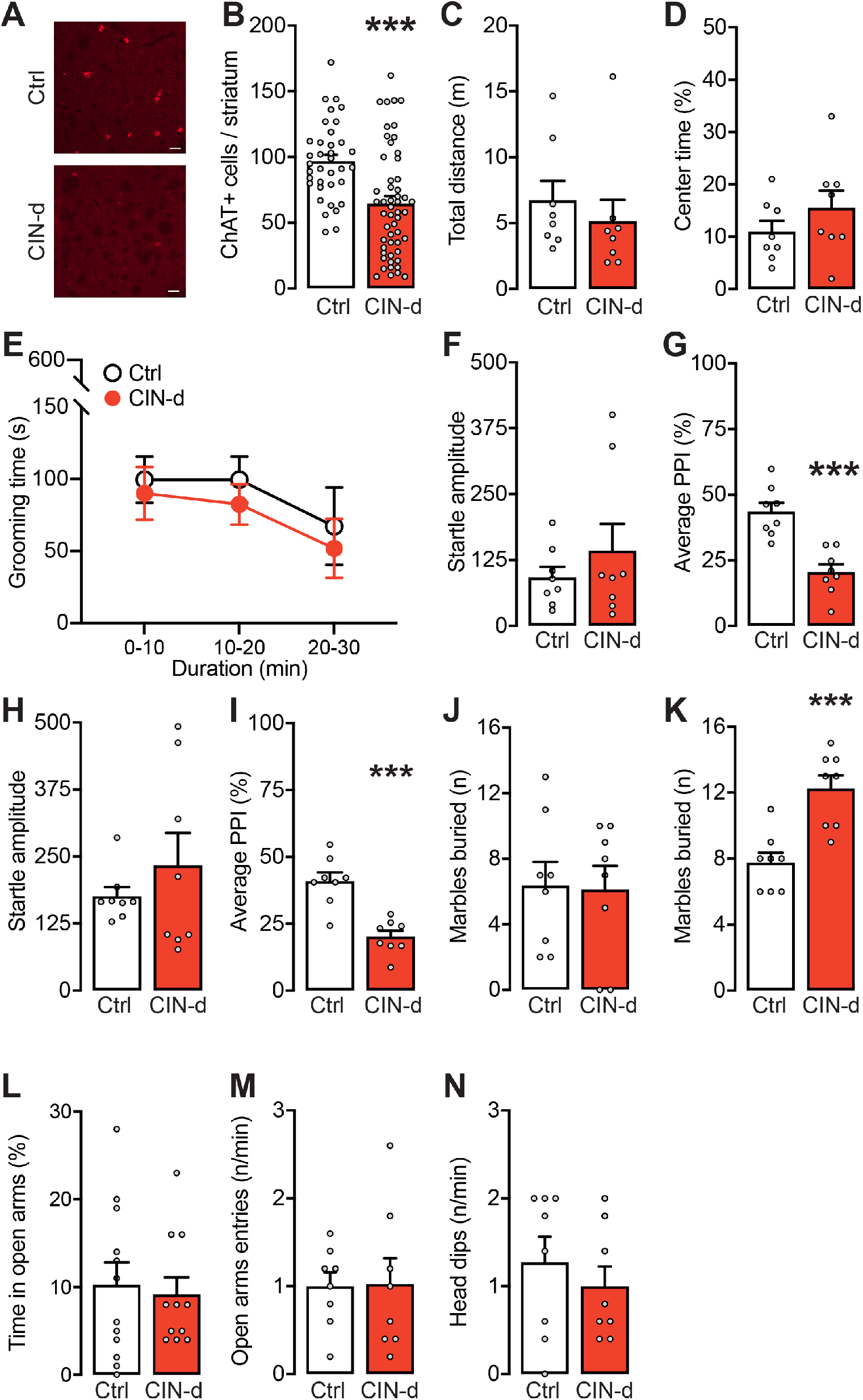
Generation and characterization of the behavioral effects of early-life CIN depletion in male mice. A-B) CIN-depletion (CIN-d) procedure led to a significant reduction of cholinergic cells in the striatum, as compared to A06-treated controls (Ctrl). Scalebars equal 10 μm. C-N) The characterization of behavioral responses of CIN-d and Ctrl mice revealed equivalent responses in C-D) open-field locomotion, E) selfgrooming behavior, and F) acoustic startle reflex; however, G) CIN-d mice exhibited significant PPI deficits. While the two groups of mice showed H) similar tactile startle amplitude, I) tactile-acoustic PPI was significantly impaired in CIN-d mice. Marbleburying behavior in CIN-depleted mice was J) equivalent to that of Ctrl animals under normal environmental light but K) significantly higher than Ctrl mice under bright light. LN) No differences between the two groups were found in the elevated plus maze. Values are presented as means ± SEM. ***, P<0.001 in comparisons with Ctrl mice.

Behavioral testing of CIN-depleted and control male mice revealed no baseline differences in open-field locomotor activity (Fig. 1C-D) or spontaneous grooming (Fig. 1E). To investigate whether early-life CIN depletion alters sensorimotor gating, male mice were tested for prepulse inhibition (PPI) of the acoustic startle. CIN-depleted mice showed no alterations in startle amplitude (Fig. 1F) but a marked reduction in acoustic PPI (Fig. 1G). Similar deficits were found in cross-modal PPI, using acoustic prepulses before tactile pulses (Fig. 1H-I). To test whether CIN depletion alters other behavioral measures associated with compulsivity, we tested male CIN-depleted mice using the marble-burying task. CIN-depleted (A06-treated) and control (C06-treated) males exhibited equivalent responses when this test was conducted under regular environmental light (Fig. 1J). However, exposure to bright light – a well-established anxiogenic condition for mice – elicited a significant enhancement in marble-burying behaviors in CIN-depleted males compared to controls (Fig. 1K). To verify whether this differential response reflected different levels of anxiety, we tested CIN-depleted mice in the elevated plus maze. However, this task revealed no significant differences with control animals (Fig. 1L-N).

Prompted by these results, we investigated whether exposure to a mild stressor that increases brain AP levels may worsen TS-related behaviors. To this end, we used spatial confinement, a manipulation shown to exacerbate tic-like responses in another model system^26^. Spatial confinement elicited an increase in grooming (Fig. 2A) and digging in CIN-depleted males (Fig. 2B). Spatial confinement also revealed increases in other sudden, fine motor responses, including axial head jerks (Fig. 2C) and eye blinks (Fig. 2D). Finally, under spatial confinement conditions, CIN-depleted mice exhibited a led to a significant increase in startle amplitude (Fig. 2E) and PPI deficits in comparison with their controls (Fig. 2F).

**Fig. 2.**
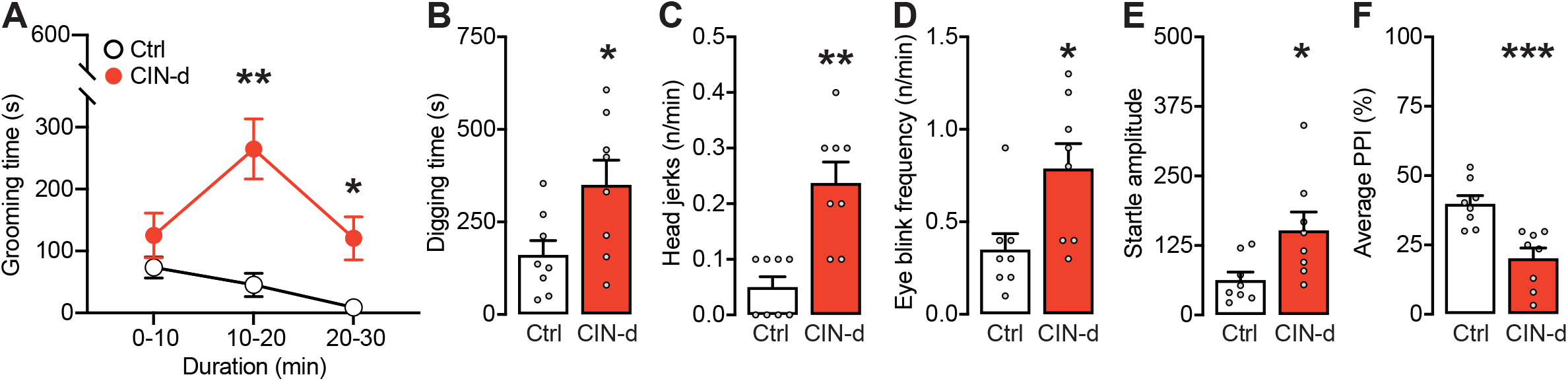
Effects of spatial confinement on the behavior of CIN-depleted (CIN-d) and control (Ctrl) male mice. Spatial confinement significantly increased A) grooming and B) digging stereotypies, as well as C) head jerks and D) eye blinks. Spatial confinement stress also E) increased acoustic startle amplitude and F) reduced PPI. Values are presented as means ± SEM. *, P<0.05; **, P<0.01; ***, P<0.001 in comparisons with Ctrl mice.

### Early-life CIN depletion fails to elicit TS-related behavioral alterations in female mice

We next tested the effects of CIN depletion in females. After verifying that the CIN depletion procedure in the striatum was successful (Fig. S1A-B), we repeated the same behavioral assessment performed in males. Most spontaneous behavioral responses were similar to controls (Fig. S1C-K), except for a significant increase in startle amplitude in CIN-depleted females (Fig. S1F). Spatial confinement failed to induce any behavioral differences between CIN-depleted and control females (S1L-N), except for higher startle amplitude (Fig. S1M).

### Tic-like behaviors are sensitive to benchmark TS therapies in CIN-depleted mice

To verify the predictive validity of the observed TS-like behavioral alterations in CIN-depleted mice, we tested the effects of several categories of drugs known to reduce the severity of tics and TS-associated symptoms. Testing was performed in CIN-depleted male mice under conditions of spatial confinement. The potent D_1_ receptor antagonist SCH23390 (0.25-0.50□mg/kg, IP, injected 15 min before testing) reduced stress-elicited increases in grooming (Fig. 3A) and digging (Fig. S2A) and rescued the PPI disruption, without changing startle responses (Fig. 3B-C). Similar ameliorative effects were elicited by clonidine (10□μg/kg, IP, injected 15 min before testing, Fig. 3D-F) and the D_2_ receptor antagonist haloperidol (0.25-0.50□mg/kg, IP, injected 15 min before testing) (Fig.3G-I and Fig. S2B). Haloperidol also rescued other fine motor manifestations, including sporadic head jerks (Fig. 3J) and eye blinks (Fig. 3K).

**Fig. 3.**
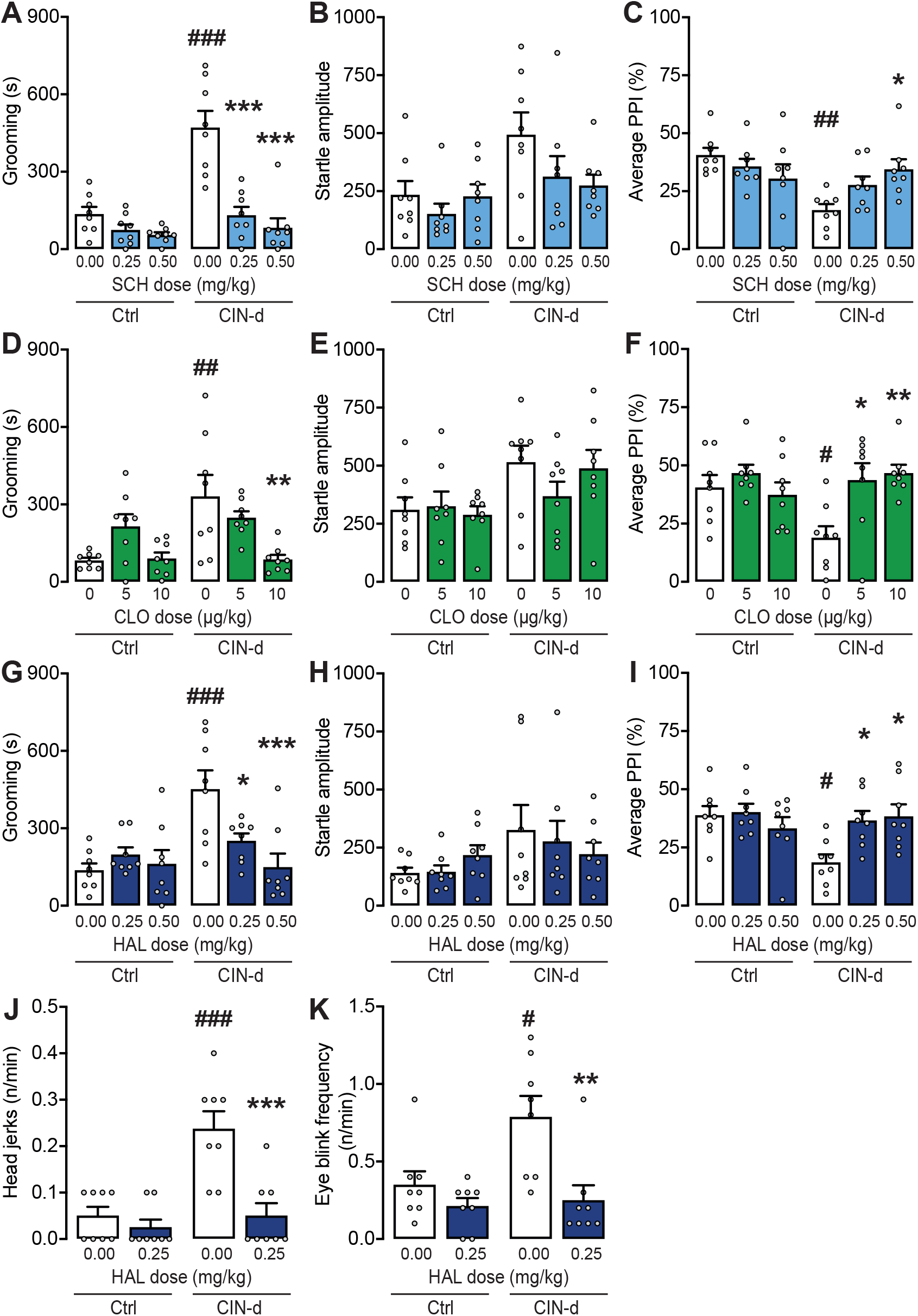
Behavioral effects of A-C) SCH 23390 (SCH), D-F) clonidine (CLO), and G-K) haloperidol (HAL) on CIN-depleted (CIN-d) and control (Ctrl) mice under spatial confinement. Values are presented as means ± SEM. ^#^, P<0.05; ^##^, P<0.01; ^###^, P<0.001 in comparisons with stressed, vehicle-treated Ctrl mice. *, P<0.05; **, P<0.01; ***, P<0.001 in comparison with stressed vehicle-treated CIN-d mice.

### AP elicits and worsens TS-related responses in freely moving CIN-depleted mice

We then tested whether AP can produce or exacerbate TS-relevant behaviors in freely moving CIN-depleted male mice in comparison with their controls. While systemic AP injections (6-12 mg/kg, IP, 5 min before testing) did not alter grooming behavior in control mice (Fig. 4A), they dose- and time-dependently increased grooming in CIN-depleted mice (Fig. 4B). We then tested the effects of AP on PPI at 6 mg/kg (IP) - a low systemic dose that does not intrinsically impair sensorimotor gating in control mice^37^. As expected, AP elicited no significant effect on startle amplitude in either control or CIN-depleted mice (Fig. 4C). While this dose of AP did not reduce PPI in control mice, it did produce a significant additional PPI deficit in CIN-depleted animals (Fig. 4D).

**Fig. 4.**
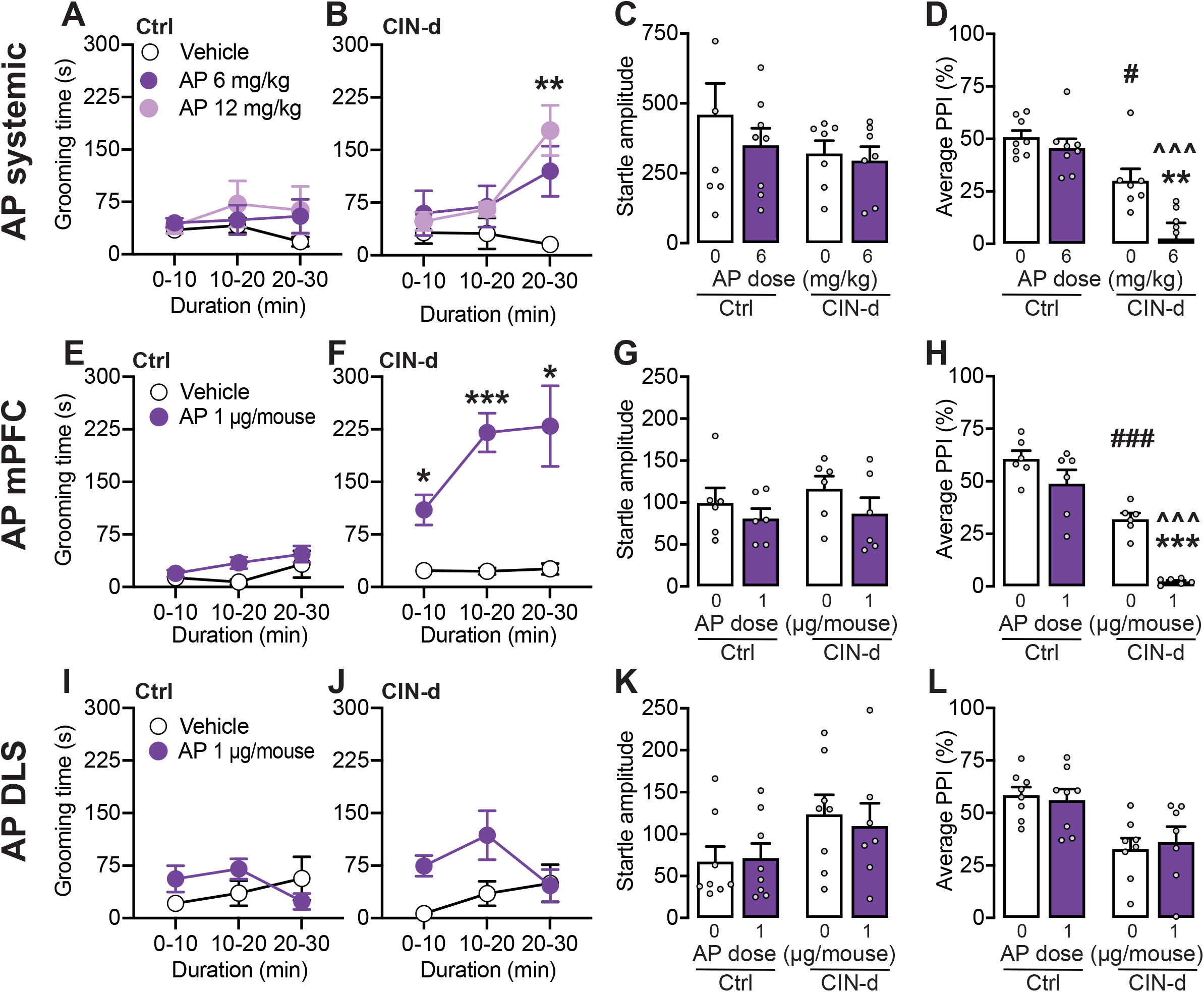
Behavioral effects of allopregnanolone (AP) on freely moving CIN-depleted (CIN-d) and control (Ctrl) mice. A-D) Effects of systemic administration; E-H) Effects of intra-medial prefrontal cortex (mPFC) infusion; I-L) Effects of intra-dorsal lateral striatum (DLS) administration. Values are presented as means ± SEM. *, P<0.05; **, P<0.01; ***, P<0.001 in comparisons with vehicle-treated Ctrl mice. ^##^, P<0.01; ^###^, P<0.001 in comparison with vehicle-treated CIN-depleted mice. ^^^, P<0.001 in comparison with AP-treated Ctrl mice.

To ascertain the neuroanatomical basis of these effects, we tested the impact of local AP injections (1μg/μl/mouse) either in the medial (m)PFC or the dorsolateral striatum (DLS), two areas implicated in tic pathophysiology. In CIN-depleted males, but not in controls, cortical AP administration significantly increased time spent grooming (Fig. 4E-F) and exacerbated the basal PPI deficit (Fig. 4H), without affecting startle amplitude (Fig. 4G). In contrast, intra-DLS administration of AP did not elicit any significant behavioral effects (Fig. 4I-L).

### Opposing AP synthesis and signaling reduces the effects of spatial confinement stress in CIN-depleted mice

To verify whether AP is necessary for the greater sensitivity of CIN-depleted mice to acute stress, we tested whether the AP synthesis inhibitor finasteride (FIN, 6-12 mg/kg, IP, 15 min before testing) could block the effects of spatial confinement. As expected, FIN dose-dependently blocked the stress-induced increase in grooming (Fig. 5A) and digging (Fig. S3A) and reversed PPI deficits (Fig. 5C) in CIN-depleted mice, without affecting startle amplitude (Fig. 5B). Similarly, the AP antagonist isoAP blocked the effects of spatial confinement on grooming and reversed the deficit in PPI in CIN-depleted mice when delivered systemically (10-20 mg/kg, IP, 15 min before testing; Fig. 5D-F; Fig. S3B) or directly into the mPFC (1 μg/mouse; Fig. 5G-I).

**Fig. 5.**
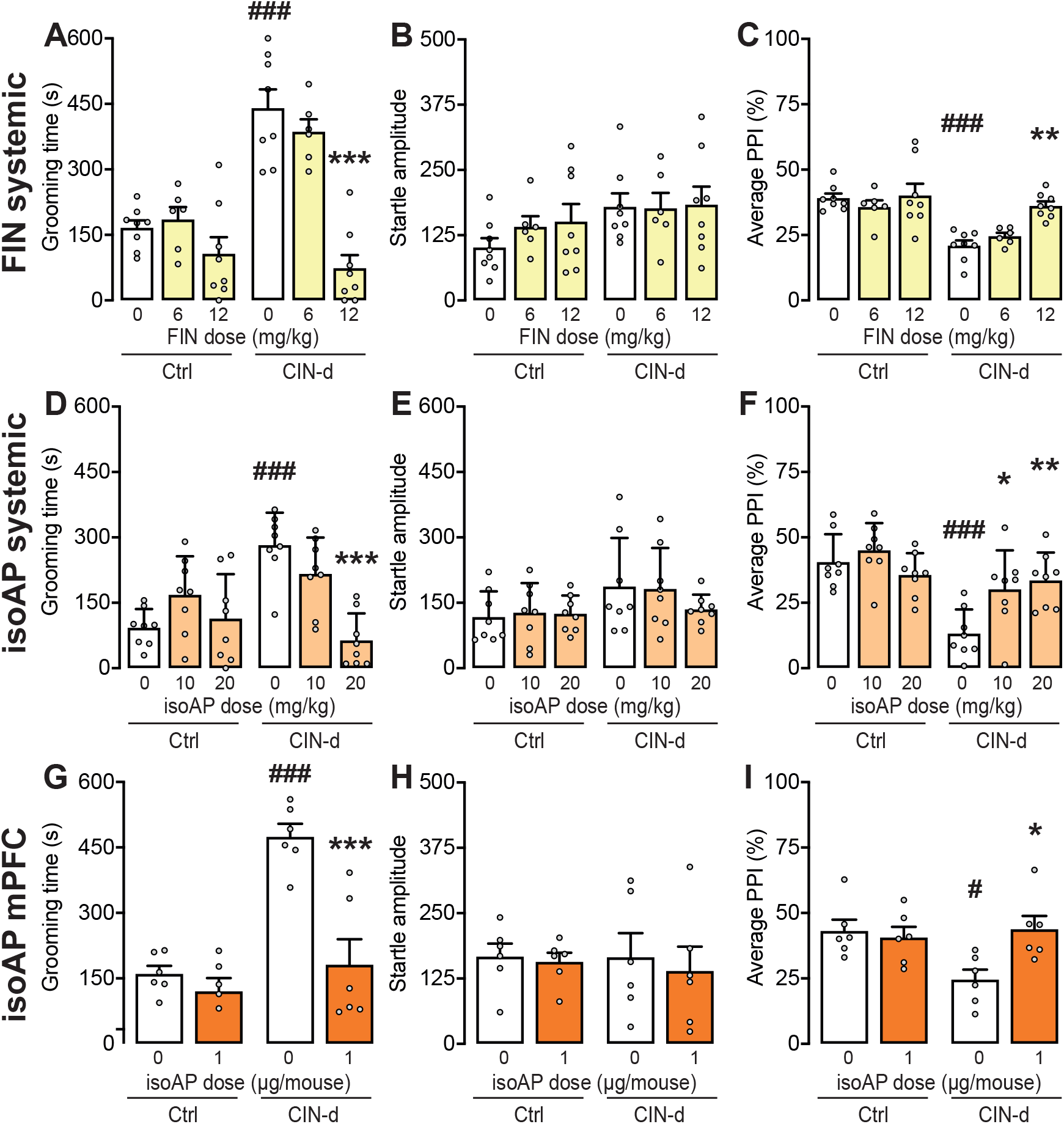
Behavioral effects of A-C) systemic finasteride (FIN), D-F) systemic iso-allopregnanolone (isoAP), and G-I) intra medial-prefrontal cortex (mPFC) isoAP on CIN-depleted (CIN-d) and control (Ctrl) mice under spatial confinement. Values are presented as means ± SEM. ^#^, P<0.05; ^###^, P<0.001 in comparisons with stressed, vehicle-treated Ctrl mice. *, P<0.05; **, P<0.01; ***, P<0.001 in comparison with stressed vehicle-treated CIN-d mice.

## DISCUSSION

In the present study, we developed a model of early-life CIN depletion using the strategy previously applied by Xu *et al*.^20^ to generate a targeted loss of dorsal striatal CINs in adult mice. In males, early-life CIN depletion resulted in a baseline deficit in PPI, which is seen in TS^38,39^. Although CIN-depleted mice did not display any alterations in baseline self-grooming and digging activity, they showed a marked increase in these stereotypies in response to spatial confinement – a mild acute stressor that had no overt effects in control mice^26^. CIN-depleted mice also exhibited elevated marble-burying, a reflection of perseverative digging^40^, but only under bright light, a well-documented environmental stressor in laboratory rodents^41^. Although grooming and digging stereotypies are typical responses to environmental stress in experimental animals^42,43^, CIN-depleted mice did not show any anxiety-like behavior in the elevated plus maze, suggesting that their stereotyped responses may reflect a greater propensity to enact repetitive behaviors in response to stress, rather than a higher sensitivity to stress itself. This interpretation is supported by previous work suggesting that striatal CIN depletion impairs behavioral flexibility. For example, several authors documented that inactivation of striatal CINs or disruption of their connectivity impairs reversal and set-shifting, but not baseline, learning^44–47^. Furthermore, striatal CIN depletion produces perseverative social responses^48^, suggesting that striatal CINs enable the extinction of adaptive and salience-driven motor sequences.

The finding of stress-sensitive stereotypies in CIN-depleted mice supports the contention that these motor responses recapitulate pathophysiological processes relevant to TS. Indeed, stereotypies share both neurobiological mechanisms and phenomenological characteristics with tics, including their sensitivity to contextual stressors^49^. From this perspective, our findings highlight the model of early-life CIN depletion as a tool with high face and construct validity, optimally suited to study the neurobiological foundations of TS and comorbid disorders. Notably, the value of these models with respect to TS is further supported by the sexual dimorphism of its behavioral manifestations, which parallels the male preponderance of TS^50^. Furthermore, the high predictive validity for TS was confirmed by our finding that the behavioral abnormalities featured by CIN-depleted mice were reversed by haloperidol and clonidine, two benchmark therapies for tic disorders, and D_1_ dopamine receptor antagonists, which have been recently highlighted as a possible new family of treatments for TS^51^. These results indicate that CIN-depleted mice may also serve as a useful platform to screen new therapeutics for tic disorders.

We have further shown that the effects of an acute mild stressor on CIN-depleted mice depend on the neurosteroid AP in the PFC. Systemic and intra-PFC administration of AP elicited behavioral responses akin to those of acute stress in CIN-depleted mice, and blocking AP synthesis (via FIN) or signaling (via isoAP) reversed the effects of stress in CIN-depleted mice without altering the behavior of control mice. These findings confirm and extend previous evidence on the key role of AP in mediating the adverse behavioral effects on TS-relevant behaviors in D1CT-7 mice, a classical model of TS with high face and predictive validity^27^. Furthermore, our data align strongly with previous results on the efficacy of FIN and isoAP in reducing stereotyped behaviors and PPI deficits across several animal models, including D1CT-7 mice^31,52,53,30,37^. Notably, the effects of FIN and isoAP were comparable to those elicited by haloperidol, suggesting that the negative modulation of AP functions may elicit similar therapeutic effects. In line with this idea, we previously showed, in a proof-of-concept open-label trial, that FIN reduced the severity and frequency of tics and compulsions in adult male TS patients^54^. A placebo-controlled trial of isoAP in TS is ongoing (Clinical Trial NCT05434546).

We previously showed that spatial confinement and other acute stressors increases AP levels in the PFC in intact wildtype mice^27,37^. The best-defined mechanism of action of AP is the positive allosteric modulation of GABA_A_ receptors^28,55,56^ through activation of a specific site in the α subunit of these channels^57,58^. At higher (micromolar) concentrations, AP acts as a GABA_A_ receptor agonist by binding to a second site located at the interface between α and ß subunits^58^. In line with these mechanisms, the detrimental effects of AP on PPI were opposed by isoAP, a selective AP antagonist that opposes its GABAergic mechanisms ^59,60^.

Ample evidence shows that the PFC sends direct efferent projections to the striatum^61^, which subserve inhibitory behavioral control^62^. Local GABAergic inhibition of projection cells in the mPFC may disinhibit perseverative responses in the context of other local striatal predisposing factors, such as a partial loss of CINs. Given that AP activates both extrasynaptic and synaptic GABA_A_ receptors, and that its affinity for different targets is influenced by the specific subunit composition of these ionotropic channels^63^, it is likely that sudden elevations of AP in the mPFC can alter the balance between excitatory and inhibitory neural activity in this region and in its connectivity with the striatum. From a translational perspective, it is worth noting that activation of the PFC has been extensively linked to tic suppression in TS^64–66^, and that preliminary evidence suggests that the main adverse effect of acute stress in TS may lie in the disruption of tic suppression efforts^67^ (see ^68^ for a detailed discussion of this potential mechanism).

The observation of spatially confined mice allowed for the characterization of other spontaneous, sporadic fine movements, such as eyeblink and head jerks, that could not be monitored in freely moving conditions. Notably, CIN-depleted mice exhibited higher frequencies of both of these responses in comparison with controls. Given that both motor manifestations bear striking isomorphism to simple tics and are reduced by haloperidol, it is tempting to speculate that these behaviors may reflect an increase in TS-related motor responses. However, given that these responses could only be captured under spatial confinement, it remains unclear whether AP plays a role in these alterations.

In contrast with adult mice that underwent dorsolateral striatal CIN depletion in adulthood^20^, our juvenile CIN depletion model showed spontaneous PPI deficits in males and increased startle response in females. This discrepancy may reflect a greater sensitivity of younger mice to the impact of CIN depletion on the regulation of sensorimotor gating; alternatively, the observed PPI deficits may reflect a partial loss of CINs in the ventral striatum (which plays a direct role in the organization of PPI) due to spread of the A06-harboring viral vector at postnatal day 4. It is possible that this effect was detected only in females due to their lower baseline levels of startle^69^.

Several limitations of the study should be recognized. First, although our results point to a key role of AP in the regulation of TS-like responses in CIN-depleted mice, we cannot exclude the possibility that other neurosteroids also participate in the adverse behavioral effects of stress. Further investigations will need to focus on other neurosteroids increased by stress, such as tetrahydrodeoxycorticosterone, or androgenic neuroactive steroids, as well as the role of the GABA_A_ receptor and other AP-sensitive receptors. Second, while our results indicate that AP mediates its effects through activation of GABA_A_ receptors in the PFC, the specific downstream mechanisms whereby this process interferes with tic execution remain unclear. Despite these caveats, our study provides support for the hypothesis that AP plays an important role in the modulation of TS-related phenotypes, especially in the context of stress, and that isoAP and FIN may be useful therapeutic tools for the reduction of tics. Given the complementary value of CIN-depleted and D1CT-7 mice in capturing different aspects of TS phenomenology and pathophysiology^49,70^, our convergent results in these two studies^27^ provide synergistic support for this mechanism in the regulation of tic fluctuations in TS. Future studies will be essential to confirm these findings in TS patients and to further explore the therapeutic potential of neurosteroid-targeting therapies in tic disorders.

## Acknowledgments

This study was supported by the NIH grants R21 NS108722 (to MB and CP) and R21 NS125654 (to MB). We would like to thank Dr. Caterina Branca for her precious editorial assistance and suggestions.

## SUPPLEMENTARY FIGURE LEGENDS

**Fig. S1. Generation and characterization of the behavioral effects of early-life CIN depletion in female mice.** A-B) CIN-depletion (CIN-d) procedure led to a significant reduction of cholinergic cells in the striatum, as compared to controls (Ctrl). Scalebars equal 10 μm. C-N) The characterization of behavioral responses of CIN-d and Ctrl mice revealed equivalent responses in C-D) open-field locomotion and E) self-grooming behavior. While CIN-depleted females displayed F) higher amplitude of the acoustic startle reflex than Ctrl, G) both groups exhibited similar PPI levels. Marble-burying behavior in CIN-depleted mice was equivalent to that of Ctrl animals in conditions of either H) normal or I) bright environmental light. J-L) No differences between the two groups were found in the elevated plus maze. Under spatial confinement, CIN-depleted had no significant effects on M) grooming behavior. CIN-depleted females exhibited N) significant enhancements of startle reflex, but O) no PPI alterations. Values are presented as means ± SEM. *, P<0.05; ***, P<0.001 in comparisons with Ctrl mice.

**Fig. S2. Behavioral effects of A) SCH 23390 (SCH) and B) haloperidol (HAL) on CIN-depleted (CIN-d) and control (Ctrl) mice under spatial confinement.** Values are presented as means ± SEM. ^#^, P<0.05; ^###^, P<0.001 in comparisons with stressed, vehicle-treated Ctrl mice. **, P<0.01; ***, P<0.001 in comparison with stressed vehicle-treated CIN-d mice.

**Fig. S3. Behavioral effects of A) finasteride (FIN) and B) iso-allopregnanolone (isoAP) on CIN-depleted (CIN-d) and control (Ctrl) mice under spatial confinement.** Values are presented as means ± SEM. ^##^, P<0.01; ^###^, P<0.001 in comparisons with stressed, vehicle-treated Ctrl mice. *, P<0.05; ***, P<0.001 in comparison with stressed vehicle-treated CIN-d mice.

