## Supplementary figures and images for "Prefrontal allopregnanolone mediates the adverse effects of acute stress in a mouse model of tic pathophysiology"

### SUPPLEMENTAL FIGURE-1

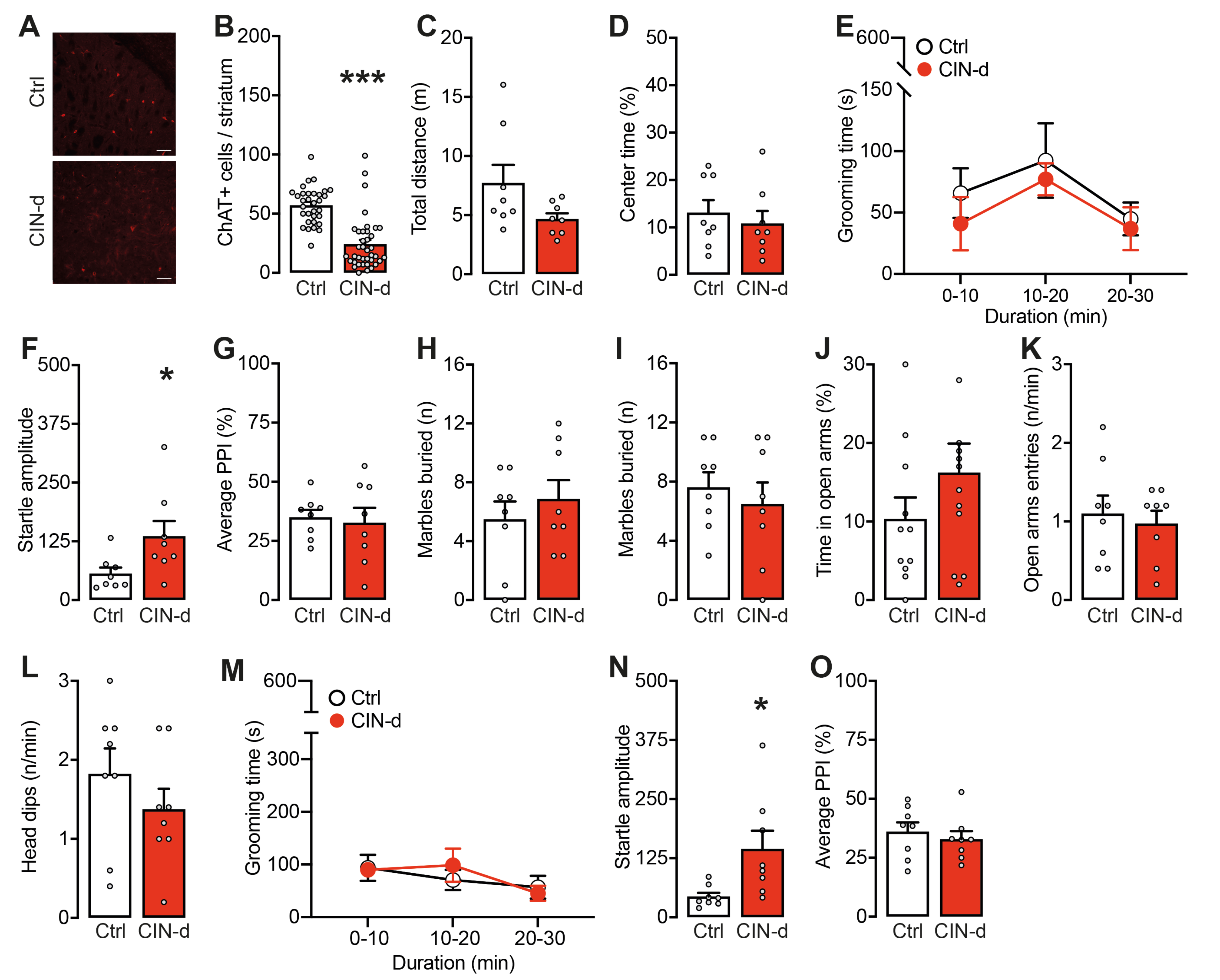

### SUPPLEMENTAL FIGURE-2

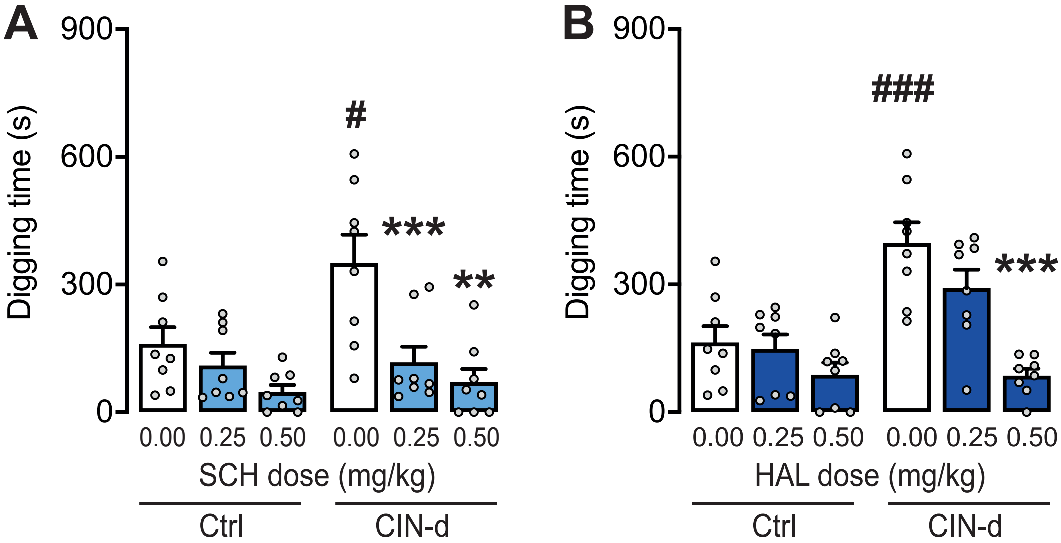

### SUPPLEMENTAL FIGURE-3

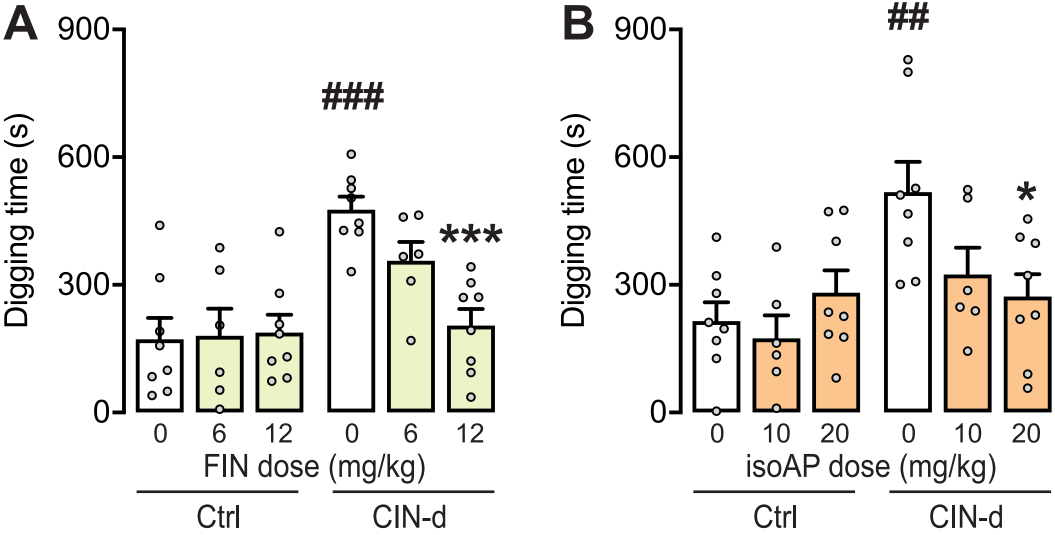
